# Effect of *Lactobacillus* postbiotic supplementation on intestinal permeability, nutrient digestibility, and gut microbial and metabolic profiles in aging cats

**DOI:** 10.64898/2026.11.09.764842

**Authors:** Sergio Andreu-Sánchez, Balint Tamasi, Anne-Sophie Boucard, Elena Pagani, Marco A. van den Berg, Brian Klein, Daniel R. Bernstein, Margreet Hekman, David G. Thomas, Karl Fraser, Erik Eckhardt

**Author notes:** Correspondence: Erik Eckhardt.

## Abstract

Aging in cats is associated with declining nutrient digestibility and increased intestinal permeability, but evidence supporting microbiome-targeted interventions in older cats remains limited. We investigated whether daily supplementation with the Lactobacillus-derived postbiotic LB fermentate HF (LBHF) affected intestinal permeability, apparent total tract digestibility, gut microbiome composition, and fecal and plasma metabolomic profiles in older cats. Twenty cats aged 7.7–13.8 years were enrolled in a blinded, placebo-controlled crossover trial comprising two 29-day treatment periods separated by a 28-day washout; 18 cats completed both periods. Intestinal permeability was the primary endpoint, while nutrient digestibility, microbiome composition, and metabolomic profiles were secondary endpoints. LBHF did not affect intestinal permeability or absorptive capacity relative to placebo, although intestinal permeability was positively associated with age. Overall nutrient digestibility was higher during LBHF treatment, with the clearest endpoint-specific evidence observed for protein digestibility. LBHF was associated with a non-significant overall increase in Shannon diversity and a reduction in the relative abundance of *Streptococcus lutetiensis*. The increase in Shannon diversity was significant only in cats receiving placebo followed by LBHF, indicating a treatment-sequence-dependent response. Thirty-three plasma and 29 fecal metabolites showed significantly different changes during LBHF treatment compared with placebo. Plasma metabolites generally showed more positive changes during LBHF treatment, whereas most associated fecal metabolites showed more negative changes. Sequence-stratified microbiome and metabolome responses differed substantially, suggesting possible period, treatment-order, or carryover effects. Overall, LBHF did not improve intestinal permeability but was well tolerated and improved nutrient digestibility, particularly protein digestibility, while producing modest and sequence-dependent microbiome and metabolome responses. These findings suggest that the effects of LBHF in healthy older cats may relate more strongly to digestive efficiency and host metabolic responses than to intestinal permeability.

## Introduction

As the lifespan of companion animals increases, maintaining quality of life in older animals may require different care strategies than for younger animals. In cats, aging is associated with physiological and nutritional changes, including loss of body weight and lean mass, reduced protein and fat digestibility, and increased intestinal permeability (Jr et al., 2008; Teshima et al., 2010; Patterson et al., 2026). The intestinal barrier permits the selective absorption of nutrients while limiting the translocation of microorganisms, toxins, and other potentially harmful substances. Disruption of this function can contribute to inflammation, food hypersensitivity, and nutrient malabsorption (Montoro-Huguet et al., 2021; Niewiem and Grzybowska-Chlebowczyk, 2022; Li et al., 2024). Gut microbial dysbiosis has been associated with increased intestinal permeability and impaired nutrient utilization. Potential mechanisms include microbiota-derived metabolites and changes in the expression or distribution of epithelial tight-junction proteins (Bischoff et al., 2014; Ghosh et al., 2021; Mishra et al., 2023; Dmytriv et al., 2024; Sandhu and Radhakrishnan, 2025).

Compared with humans and other animal species, age-related changes in intestinal barrier function, nutrient digestibility, the gut microbiome, and the metabolome remain insufficiently characterized in cats. A recent study found greater small-intestinal permeability in senior cats aged 9.5–16.2 years than in younger cats aged 1.3–3.4 years (Patterson et al., 2026). Although interactions between the gut microbiota and intestinal barrier function have been extensively studied in other species, their relationship in cats remains poorly understood, as do the mechanisms linking nutrition, the gut microbiota, and age-related changes in intestinal permeability (Laval et al., 2015; Di Vincenzo et al., 2024).

Microbiome-targeted interventions, including postbiotics, may offer one approach to supporting healthy aging in cats. Postbiotics are defined as preparations of inanimate microorganisms and/or their components that confer a health benefit on the host (Salminen et al., 2021). The *Lactobacillus*-derived postbiotic LB fermentate HF (LBHF) has been shown to strengthen intestinal epithelial integrity in human colonic organoids and to reduce *Escherichia coli*-induced epithelial stress *in vitro* and *in vivo* (Liévin-Le Moal et al., 2002; Xu et al., 2022; Duarte et al., 2024; Cercamondi et al., 2025). In adult dogs, LBHF supplementation has also been associated with changes in acute stress responses and temporal variation in the gut microbiome (Eckhardt et al., 2026). In another study in adult dogs, LBHF affected fecal microbial populations and selected immune- and stress-related outcomes (Koziol et al., 2023).

We therefore evaluated whether daily LBHF supplementation affected intestinal permeability in older cats in a randomized, blinded, placebo-controlled crossover trial. Secondary objectives were to assess its effects on apparent total tract digestibility, gut microbiome composition, and fecal and plasma metabolomic profiles.

## Materials and methods

### Cat cohort and housing

Twenty domestic shorthair cats (*Felis catus*), ranging in age from 7.7 to 13.8 years, were enrolled and assigned by stratified randomization to one of two treatment sequences: placebo followed by LBHF (placebo→LBHF) or LBHF followed by placebo (LBHF→placebo). Randomization was performed to balance age, sex, body weight, and behavioral compatibility between sequences.

Because of behavioral compatibility requirements, cats were co-housed with non-treatment conspecifics in four colony pens (1400 × 2400 × 1400 cm) throughout the study. During the apparent total tract digestibility (ATTD) total fecal collection phase, cats were temporarily housed individually in pens measuring 80 × 80 × 110 cm. Water was provided *ad libitum*, and body weight was measured weekly.

One cat died early in the study from causes unrelated to the study, and a second cat was withdrawn after being diagnosed with feline infectious peritonitis. Therefore, 18 cats completed both periods of the crossover trial, with nine cats in each treatment sequence. The sequence-level baseline characteristics shown in **Table 1** describe these 18 completers.

**Table 1.**
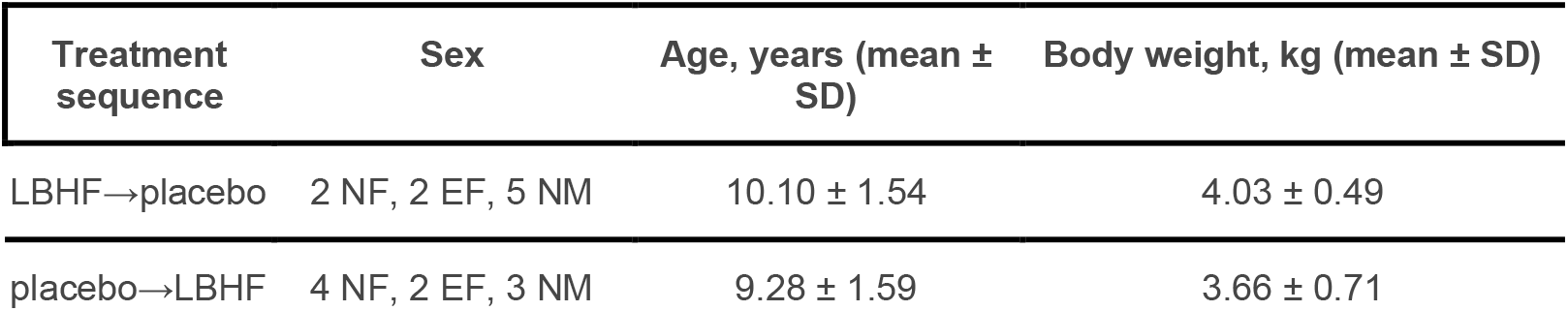
Baseline characteristics of cats that completed both treatment periods, by randomized treatment sequence (n = 9 per sequence). Values for age and body weight are presented as mean ± SD. NF, neutered female; EF, entire female; NM, neutered male.

| **Treatment sequence** | **Sex** | **Age, years (mean ± SD)** | **Body weight, kg (mean ± SD)** |
|---|---|---|---|
| LBHF→placebo | 2 NF, 2 EF, 5 NM | 10.10 ± 1.54 | 4.03 ± 0.49 |
| placebo→LBHF | 4 NF, 2 EF, 3 NM | 9.28 ± 1.59 | 3.66 ± 0.71 |

### Diet and interventions

Throughout the trial, cats were fed their usual commercially available wet food diet (Chef; Kraft-Heinz-Watties, Hastings, New Zealand). The two dietary supplement interventions, provided by dsm-firmenich, were a placebo consisting of microcrystalline cellulose and a dried *Lactobacillus* postbiotic preparation (LBHF). During each treatment period, the assigned supplement was administered orally once daily at 20 mg/kg body weight.

Each morning, cats were placed individually in a pen for approximately 10–20 min and offered the assigned supplement mixed with a small portion of their regular diet. After complete consumption of the supplemented portion, cats were returned to their respective colony pens.

Feliway® spray (synthetic feline facial pheromone analogue; Ceva Animal, Libourne, France) was applied in the individual housing environment to reduce stress-related behaviors, improve environmental familiarity, and limit stress-induced variation in physiological and behavioral responses.

### Trial design

The study used a blinded, placebo-controlled, two-period crossover design. Before the first treatment period, cats underwent a 28-day run-in period. During the first 29-day treatment period (Period A), cats received either LBHF or placebo orally once daily according to their randomized treatment sequence. This was followed by a 28-day washout period during which no intervention was administered. Cats then crossed over to the alternate intervention for a second 29-day treatment period (Period B). Thus, cats assigned to the placebo→LBHF sequence received placebo during Period A and LBHF during Period B, whereas cats assigned to the LBHF→placebo sequence received LBHF during Period A and placebo during Period B. Cats remained in colony housing except during the individual housing periods required for apparent total tract digestibility and intestinal permeability assessments, as described below.

### Intestinal permeability sampling

A sugar absorption test (SAT) was performed at the beginning (Day 0) and end (Day 29) of each treatment period. Before the baseline SAT, cats were acclimated to individual housing on Days −2 and −1 and fasted overnight for at least 12 h, with *ad libitum* access to fresh water. At baseline (T0), 2 mL of blood was collected, and plasma was obtained to assess background concentrations of the sugar probes. Plasma aliquots were stored at −80 °C until analysis.

Cats were then administered an oral solution containing lactulose, rhamnose, xylose, and 3-O-methylglucose in purified water by syringe. The solution was administered at a total volume of 2 mL/kg body weight, as previously described (Patterson et al., 2024). Food and water were withheld for 3 h after administration. A second blood sample was collected 3 h after dosing (T3), and plasma aliquots were stored at −80 °C until analysis. Cats were then returned to their colony housing and provided access to food and water. The SAT was repeated on Day 29 of each treatment period using the same procedure.

### Intestinal permeability analysis

A pooled sample of all T3 h plasma samples was prepared prior to analysis to enable multi-batch monitoring for instrument performance and consistency of results. On each analysis day, this pooled quality control (QC) sample was extracted in duplicate. Each QC extract was analyzed every 15 samples, resulting in 4 × 2 analyses of the pooled QC samples in each of the four analytical batches.

Plasma concentrations of lactulose, rhamnose, xylose, and 3-O-methylglucose were quantified as described previously (Patterson et al., 2024). Briefly, 590 μL of ice-cold acetonitrile and 10 μL of internal standard mixture containing 50 μg/mL of each of four ^13^C-labelled sugars (^13^C_12_ lactulose, ^13^C_6_ 3-*O*-methylglucose (3-OMG), ^13^C_6_ rhamnose, and ^13^C_5_ xylose) were added to an aliquot of 200 μL of plasma. Samples were vortex-mixed thoroughly, incubated (60 min, -20 °C), vortexed again, and centrifuged (14,000 × g, 10 min, 4 °C). Then, 100 μL of purified water was added to each sample, before vortexing and centrifuging (14,000 × g, 10 min, 4 °C) a second time. An 800-μL aliquot of the supernatant was transferred and dried under nitrogen and finally resuspended in 100 μL of 90% acetonitrile containing 1 mM ammonium formate and subsequently analyzed by LC-MS. Extraction blanks and pooled QC samples were prepared in the same way and included in the analysis. Extraction blanks were prepared from deionized (DI) water, and the pooled QC sample was prepared by combining aliquots from all T3 h plasma samples.

Calibration standards were prepared from stock solutions of unlabeled sugars along with the stable-isotope-labeled internal standards for each compound to give a 6-point calibration set.

LC-MS was carried out using a Thermo Scientific Accela UHPLC system coupled to a Q Exactive Orbitrap mass spectrometer (MS) with electrospray ionization (ESI). Samples (5 μL) were introduced to the HPLC system via a PAL autosampler, and separation was achieved using a Phenomenex Luna NH2 column (2 × 150 mm, 3 μm) using the published solvents and gradient. The MS was operated in negative ionization mode with full scan MS1 (m/z 50–500) and parallel reaction monitoring (PRM) MS/MS for the stable-isotope labelled and unlabeled sugars. Data was analyzed using Thermo Xcalibur Quan Browser 4.7, and all analytes were quantified against their respective calibration curves.

### Apparent Total Tract Digestibility assessment

A batch of fresh fecal samples was collected within 15 minutes of defecation at the beginning of the apparent total tract digestibility (ATTD) assessment period (Day 22). Samples were divided into four 1-g aliquots, immediately snap-frozen in liquid nitrogen, and stored at −80 °C until analysis. Feces collected for ATTD assessment and representative samples of the diet were stored at −20 °C, freeze-dried, and finely ground before analysis.

ATTD was determined according to the Association of American Feed Control Officials protocol for apparent nutrient digestibility and metabolizable energy (Association of American Feed Control Officials, 2025). Diet and fecal samples were analyzed for moisture by drying to constant weight in a convection oven at 105 °C (AOAC methods 930.15 and 925.10). Ash was determined by combustion at 550 °C (AOAC 942.05). Crude protein and crude fat were determined using the LECO total combustion method (AOAC 968.06) and acid hydrolysis/Mojonnier extraction (AOAC 954.02), respectively. Gross energy was determined by bomb calorimetry, and crude fiber was determined gravimetrically (AOAC 978.10). Nitrogen-free extract was calculated by difference (Latimer and Latimer, 2023).

### Intestinal permeability statistical analysis

Individual plasma sugar concentrations were quantified, and the lactulose:rhamnose (L:R) and xylose:3-O-methylglucose (X:G) ratios were calculated as measures of intestinal permeability and absorptive capacity, respectively. For each treatment period, within-cat changes were calculated as Δ = Day 29 − Day 0. Treatment-specific change scores were compared within cats using two-sided Wilcoxon signed-rank tests, both across the full crossover population and separately within each treatment sequence. This approach accounted for the period-specific baseline by comparing within-cat changes between LBHF and placebo. Only cats with complete measurements required to calculate the paired treatment-specific changes were included in this analysis.

As a complementary analysis, associations between age and L:R and X:G ratios were evaluated using linear mixed-effects models with age and study day as fixed effects and cat as a random intercept.

#### Body weight analysis

Longitudinal changes in body weight during the treatment periods were evaluated using a linear mixed-effects model. Body weight was modeled as a function of treatment, day within treatment period, their interaction, and study period, with a random intercept for each cat. The treatment × day interaction was used to test whether changes in body weight over time differed between LBHF and placebo.

### Statistical analysis of digestibility

Digestibility endpoints, including dry matter (DM), gross energy (GE), protein, fat, and metabolizable energy (ME), were compared between LBHF and placebo within cats. Dry matter digestibility was expressed as a percentage of total fecal mass, gross energy, protein, and fat digestibility as percentages of dry matter, and metabolizable energy as kcal/kg food. For each endpoint, LBHF and placebo measurements from the same cat were compared using a two-sided Wilcoxon signed-rank test. A permutation-based quadratic-form omnibus test was used to test the global null hypothesis of no difference between LBHF and placebo across the five digestibility endpoints. For endpoint-specific inference, single-step maxT-adjusted *P* values were calculated to control the family-wise error rate across the five endpoints using the coin R package (v1.4.4) (Hothorn et al., 2008).

As a complementary analysis, the five digestibility outcomes were standardized to z-scores and subjected to principal component analysis. The first principal component (PC1) was used as a composite measure of digestibility, and PC1 scores were compared between LBHF and placebo using a two-sided paired Wilcoxon signed-rank test.

### DNA extraction, library preparation, and sequencing

DNA was extracted from fecal samples using the DNeasy PowerSoil Pro Kit (QIAGEN) according to the manufacturer’s instructions and quantified using a Qubit Flex Fluorometer with the Qubit dsDNA HS Assay Kit (Thermo Fisher Scientific). DNA libraries were prepared using the Watchmaker DNA Library Prep Kit (7K0019-1K). Extracted DNA was fragmented using a mastermix of Watchmaker Frag/AT Buffer and Frag/AT Enzyme Mix. IDT xGen UDI Primers and IDT Stubby Adapters were added to each sample followed by 7 cycles of PCR to construct the DNA libraries. The final DNA libraries were purified using CleanNGS magnetic beads (CleanNA) and eluted in nuclease-free water. Following elution, the libraries were quantified using the Qubit^TM^ fluorometer and Qubit dsDNA HS Assay Kit. Libraries were subsequently circularized and sequenced on an Element AVITI system using the Cloudbreak FreeStyle sequencing kit to generate 2 × 150-bp paired-end reads.

### Metagenomic taxonomic profiling with SingleM

Species-level taxonomic profiles were generated from raw, untrimmed metagenomic sequencing reads using SingleM v0.19.0 in *pipe* mode (Woodcroft et al., 2026). SingleM estimates taxonomic abundance by recruiting reads to conserved regions of 59 single-copy marker genes and assigning taxonomy using Genome Taxonomy Database (GTDB) reference genomes. Raw sequencing reads were analyzed without prior host-read removal as recommended for the SingleM workflow.

### Beta diversity

Species-level relative abundances were centered log-ratio (CLR) transformed (pseudocount: 1 × 10⁻⁶). Aitchison distances were calculated as Euclidean distances between CLR-transformed profiles.

PERMANOVA (vegan v2.6.6.1, adonis2) was applied with permutations restricted within individual cats to account for repeated measurements (Dixon, 2003). Marginal effects of treatment, period, and study day were assessed using by = "margin".

### Alpha diversity

Shannon diversity was calculated from species-level relative abundances using the diversity function in the vegan R package. Within each treatment period, the change in Shannon diversity (ΔShannon) was calculated as the end-of-period value minus the beginning-of-period value (P3 – P1 for Period A and P6 − P4 for Period B). For each cat, ΔShannon during LBHF treatment was compared with ΔShannon during placebo treatment using a two-sided Wilcoxon signed-rank test. Analyses were performed in the full crossover population and separately within each treatment sequence.

### Differential abundance testing for taxa

Species detected in at least 20% of samples were included in univariate differential abundance testing. Species-level relative abundances were CLR-transformed as described above. Within each treatment period, per-cat changes in CLR values (ΔCLR) were calculated as the end-of-period value minus the beginning-of-period value (P3 − P1 for Period A and P6 − P4 for Period B). For each taxon, ΔCLR during LBHF treatment was compared with ΔCLR during placebo treatment using a two-sided Wilcoxon signed-rank test. Only cats where the species was detected at least once among the six available metagenomic samples were used. Analyses were performed in the full crossover population and separately within each treatment sequence. False discovery rate (FDR) Benjamini-Hochberg correction was applied to account for multiple testing across taxa, with FDR < 0.05 considered statistically significant.

As a sensitivity analysis to assess treatment differences without potential carryover from a preceding intervention, the analysis was restricted to Period A. For each cat, ΔCLR was calculated as P3 − P1 and compared between cats receiving LBHF and those receiving placebo using a two-sided Wilcoxon rank-sum test. The same taxon inclusion criteria and multiple-testing correction were applied.

### Metabolomic measurements

Plasma and fecal metabolites were quantified using a targeted metabolomics approach based on the Biocrates MxP® Quant 500 kit (Biocrates Life Sciences AG, Innsbruck, Austria). Samples were processed according to the manufacturer’s instructions. Sample extracts, reference standards, and quality controls were analyzed using ultra-high-performance liquid chromatography (UHPLC) and flow injection analysis coupled to tandem mass spectrometry. An Agilent 1290 Series UHPLC system coupled to a 6500+ QTRAP mass spectrometer equipped with an IonDrive Turbo V electrospray ionization source (Sciex, Foster City, CA, USA) was used for analysis. Chromatographic and mass spectrometric parameters followed the manufacturer’s protocol, and data were processed using Biocrates WebIDQ software.

Metabolomic profiling was performed on fecal and plasma samples collected at P1, P3, P4, and P6. Of the 630 metabolites included in the analytical panel, 389 were detected in fecal samples and 467 in plasma samples. Metabolites detected above the limit of detection in ≥80% of samples were retained for diversity analyses, whereas those detected in ≥20% of samples were retained for univariate analyses. Semi-quantitative values below LOD were retained for analysis.

### Metabolomics diversity and beta-diversity analyses

Metabolomic alpha- and beta-diversity analyses were performed separately for fecal and plasma samples. For alpha-diversity analyses, metabolites detected above the limit of detection in ≥80% of samples were retained. Shannon diversity was calculated using the diversity function in the vegan R package based on raw abundance values. Within each treatment period, changes in metabolomic Shannon diversity (ΔShannon) were calculated as the end-of-period value minus the beginning-of-period value (P3 − P1 for Period A and P6 − P4 for Period B). For each cat, ΔShannon during LBHF treatment was compared with ΔShannon during placebo treatment using a two-sided Wilcoxon signed-rank test. Analyses were performed both in the full crossover population and separately within each treatment sequence.

For beta-diversity analyses, metabolite intensities were standardized as z-scores per metabolite across samples, and Euclidean distances were calculated between samples. PERMANOVA with 999 permutations restricted within cat was used to assess treatment-associated differences in metabolomic profiles. Analyses were performed separately for fecal and plasma metabolomes, and the proportion of variance explained by treatment was reported as PERMANOVA R².

### Univariate metabolite testing

For each metabolite, within-cat changes (Δ) were calculated as the concentration at the end of each treatment period minus the corresponding period baseline. Treatment-associated differences in Δ-values between LBHF and placebo were assessed using paired Wilcoxon signed-rank tests. Analyses were performed both overall and separately within each treatment sequence. P-values were adjusted for multiple testing using the Benjamini-Hochberg FDR, with FDR < 0.05 considered statistically significant. Effect sizes were additionally estimated using Cohen’s d. For chemical-class enrichment analysis, metabolites were ranked according to −log10(P) × sign(d), combining the strength and direction of the treatment-associated difference. Gene Set Enrichment Analysis (GSEA) was performed using fgsea v1.26.0, with minimum and maximum class sizes of 3 and 500 metabolites, respectively (Korotkevich et al., 2021).

### Univariate associations between omics features and digestibility

To identify omics features associated with nutrient digestibility, each digestibility endpoint was modeled as a function of individual features from three omics layers: species-level microbial relative abundances (CLR-transformed), fecal metabolite abundances (log-transformed), and plasma metabolite abundances (log-transformed). Digestibility measurements from each treatment period were paired with omics measurements collected at the corresponding end-of-period sampling point. For metabolites measured in both plasma and feces, plasma-to-fecal log-ratios were calculated as log(plasma) − log(feces) after matching measurements by cat and time point.

For each endpoint-feature pair, a linear mixed-effects model was fitted with the digestibility endpoint as the response; the omics feature, treatment, period, treatment sequence, age, and sex as fixed effects; and cat as a random intercept. Benjamini-Hochberg correction was applied separately within each digestibility endpoint and omics layer.

To identify chemical classes associated with digestibility endpoints, class-level enrichment analyses were performed separately for each endpoint. Metabolites were ranked according to −log10(*P*) × sign(β), where β represents the estimated association between metabolite abundance and the digestibility endpoint. GSEA was performed separately for fecal metabolites, plasma metabolites, and plasma-to-fecal log-ratios using metabolite annotations to define chemical classes.

## Results

### Intestinal permeability increased with age but did not differ between LBHF and placebo

Eighteen cats completed both periods of the crossover trial, with nine cats in each treatment sequence (**Figure 1**). The administered LBHF dose was similar between study periods (19.94 ± 0.019 mg/kg in Period A and 19.92 ± 0.019 mg/kg in Period B; *P* = 0.94). Supplement compliance was high, body weight remained stable throughout the study, and no treatment-related clinical concerns were reported.

**Figure 1.**
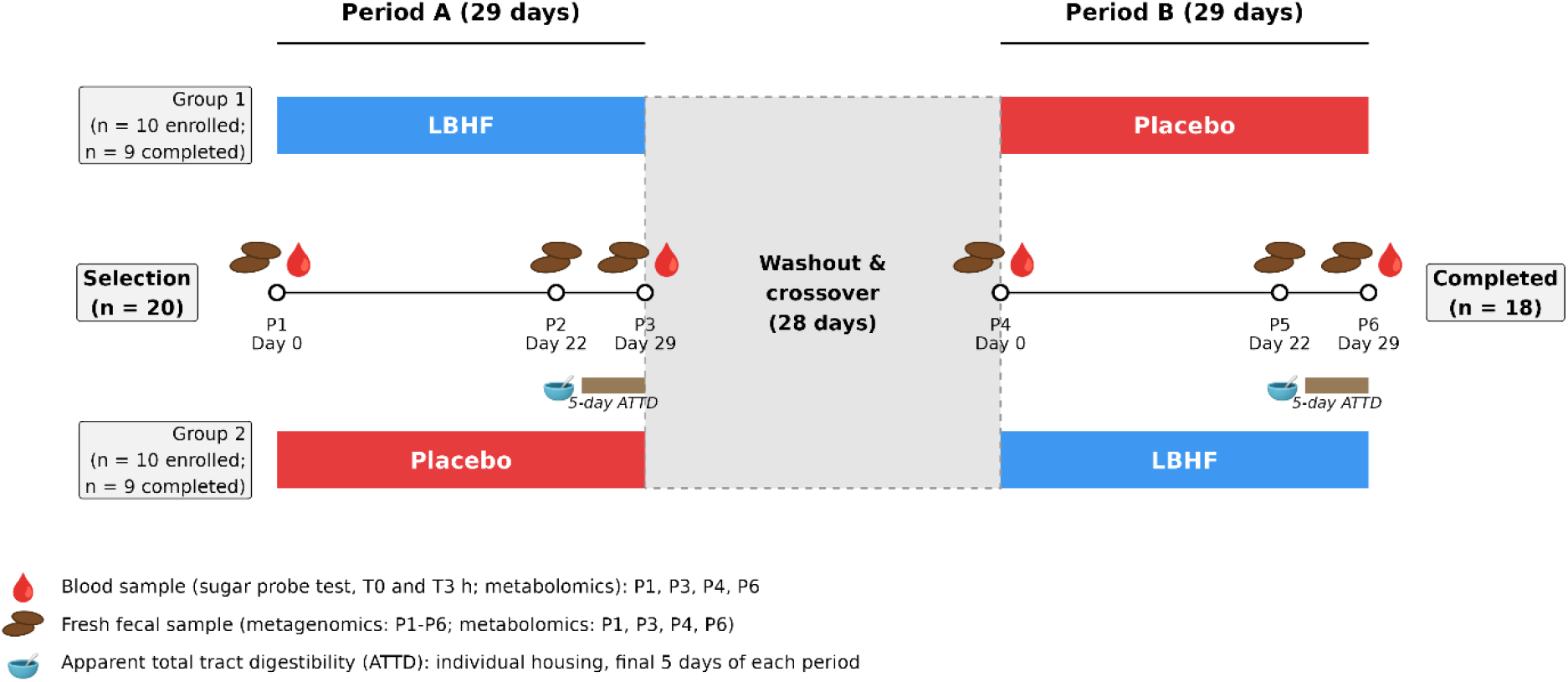
Study design. Twenty cats were assigned to one of two treatment sequences in a randomized, placebo-controlled crossover trial: LB fermentate HF (LBHF) followed by placebo, or placebo followed by LBHF. Each treatment period lasted 29 days and was separated by a 28-day washout period. Ten cats were initially assigned to each sequence; 18 cats completed both treatment periods. Cats were group housed except during the final 5 days of each treatment period, when they were individually housed for apparent total tract digestibility measurements. Fecal samples were collected during each treatment period, and blood samples for intestinal permeability assessment were collected at baseline and at the end of each treatment period.

Intestinal permeability and absorptive capacity were assessed using the lactulose:rhamnose (L:R) and xylose:3-O-methylglucose (X:G) ratios, respectively. None of the four sugar probes was detected in baseline (T0) plasma samples, and T3 concentrations were within previously reported ranges (Patterson et al., 2024). Complete measurements were available for 17 cats; one cat was excluded from the permeability analysis after failing to ingest the sugar solution at one sampling point.

Within each treatment period, changes in L:R and X:G ratios were calculated as the Day 29 value minus the Day 0 value. No evidence of a difference between LBHF and placebo was detected for either intestinal permeability (L:R; Wilcoxon signed-rank test, *P* = 0.70) or absorptive capacity (X:G; *P* = 0.46) (**Figure 2A,B**). No significant differences were detected in treatment-sequence-specific analyses. Age was positively associated with the L:R ratio (β = 0.03, *P* = 4.76 × 10⁻⁴; **Figure 2C**), whereas no association was detected between age and the X:G ratio (β = 0.01, *P* = 0.80; **Figure 2D**)

**Figure 2.**
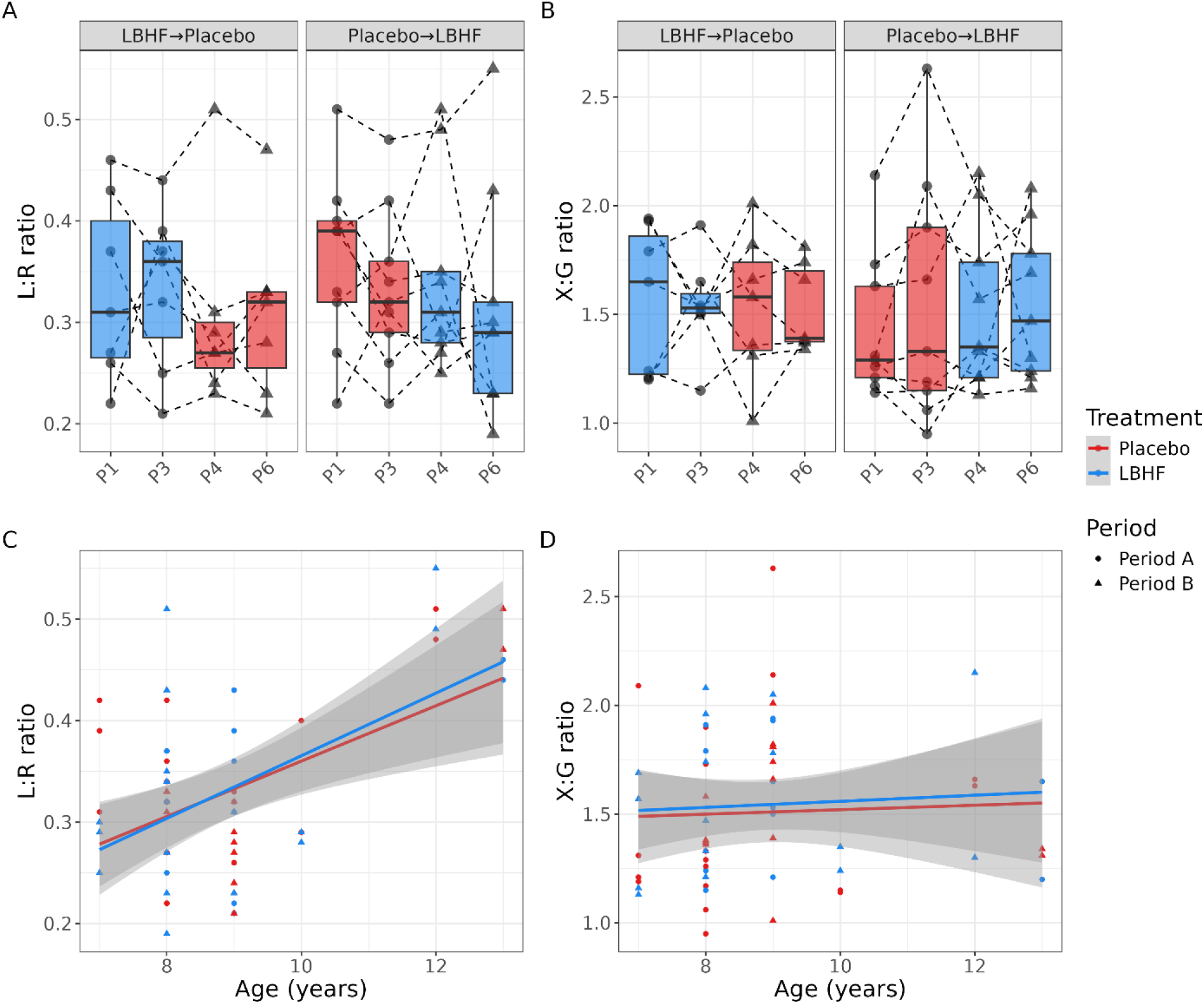
Intestinal permeability and absorptive-capacity readouts throughout the study. **(A, B)** Temporal variation in lactulose:rhamnose (L:R) ratio (A) and xylose:3-O-methylglucose (X:G) ratio (B). No evidence of treatment effect is observed in overall or per-group analysis. **(C, D)** Chronological age relationship with L:R ratio (C) and X:G ratio (D).

### LBHF supplementation was associated with improved nutrient digestibility

Throughout the study, body weight remained generally stable (**Table S1**), and the rate of change in body weight did not differ significantly between LBHF and placebo (treatment × day interaction: β = −1.87 g/day, SE = 1.27, *P* = 0.14). Together with high supplement compliance and the absence of reported treatment-related clinical concerns, these findings support that LBHF was well tolerated during the study.

Apparent total tract digestibility was consistently numerically higher during LBHF treatment than during placebo across all five evaluated endpoints (**Table 2**; **Figure 3A–E**). The global omnibus test indicated an overall difference in digestibility between treatments (*P* = 0.016). In endpoint-specific analyses, protein digestibility increased from a median of 83.0% during placebo treatment to 84.5% during LBHF treatment (Cohen’s *d* = 0.58, unadjusted *P* = 0.018, multiplicity-adjusted *P* = 0.049). None of the other individual digestibility endpoints reached statistical significance after adjustment for multiple testing (**Table 2**).

**Figure 3.**
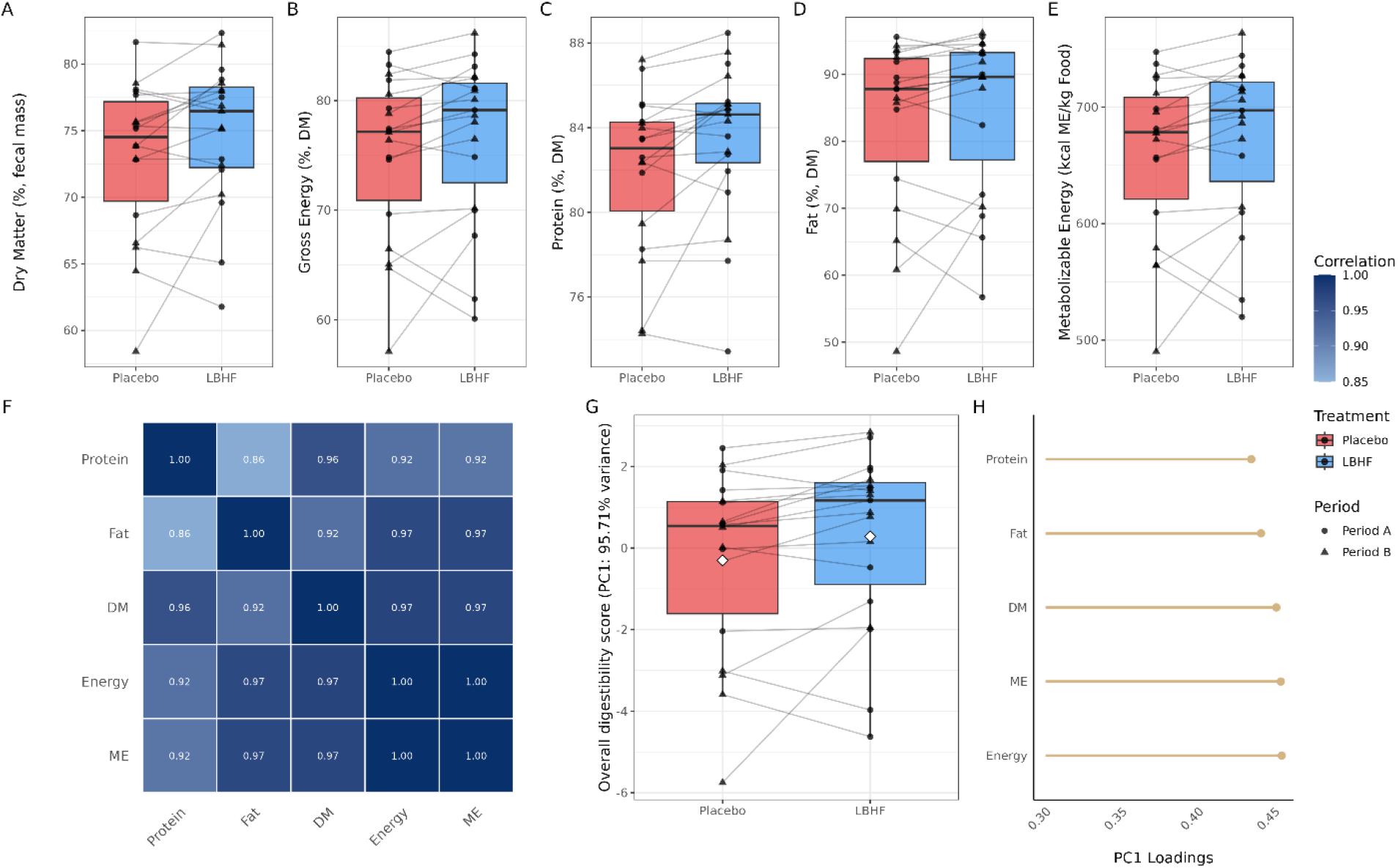
LBHF supplementation was associated with greater nutrient digestibility. **(A–E)** Apparent total tract digestibility measurements in samples treated with placebo or LBHF. (**F)** Pearson correlations between digestibility of different nutrients. **(G)** Changes of a composite score of digestibility (PC1 of digestibility features) during the cross-over study. **(H)** Loadings of PC1 highlight the weighted contribution of each nutritional feature in the composite score displayed in panel G.

**Table 2.**
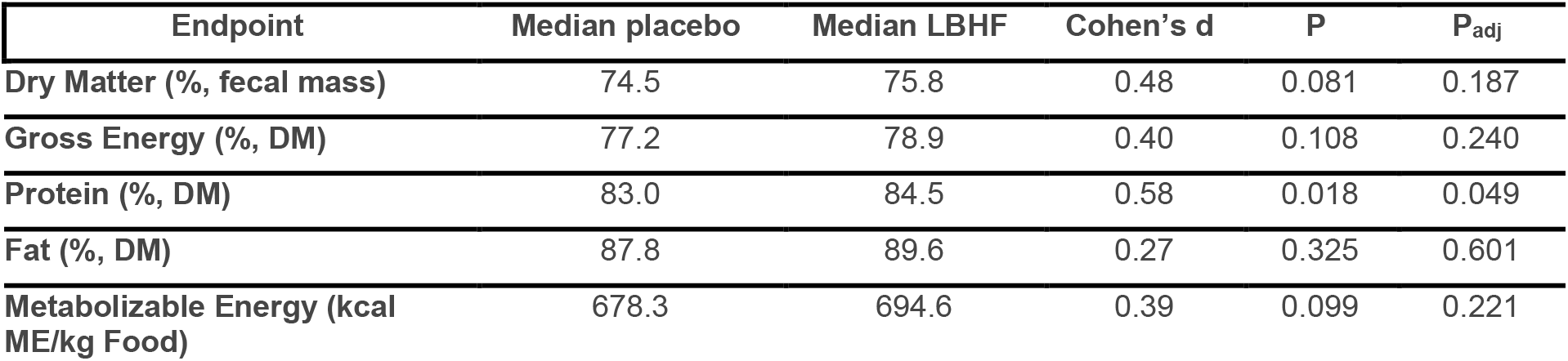
Apparent total tract digestibility for placebo and LBHF fed to n = 18 cats. Values are medians for each treatment. Cohen’s d, unadjusted P values, and multiplicity-adjusted P values from paired Wilcoxon signed-rank tests are shown.

Digestibility endpoints were strongly correlated with one another (*r* = 0.86–1.00; **Figure 3F**). Consistent with this covariance, the first principal component (PC1) of the five standardized digestibility endpoints explained 95.7% of their total variance (**Figure 3G,H**). PC1 scores were numerically higher during LBHF treatment, although the difference did not reach statistical significance (paired Wilcoxon signed-rank test, *P* = 0.06). Together, the significant global test and endpoint-specific increase in protein digestibility indicate that LBHF supplementation was associated with improved nutrient digestibility, with the clearest evidence observed for protein digestibility.

### Gut microbiome responses differed by treatment sequence

Shotgun metagenomic sequencing generated a median of 5.48 million paired-end reads per sample (range, 3.07–48.55 million). SingleM profiling identified 309 microbial species spanning 162 genera and 12 phyla across the dataset (**Supplementary Figure 1**).

We used PERMANOVA based on Aitchison distances to evaluate factors associated with gut microbial community composition. Treatment sequence, study period, and study day were associated with community composition (R² = 0.020, P = 0.010; R² = 0.013, P = 0.004; and R² = 0.010, P = 0.020, respectively). The corresponding ordination is shown in **Figure 4A**. In sequence-stratified PERMANOVAs, treatment was not significantly associated with community composition in either the Placebo→LBHF sequence (R² = 0.018, P = 0.056) or the LBHF→Placebo sequence (R² = 0.020, P = 0.065).

**Figure 4.**
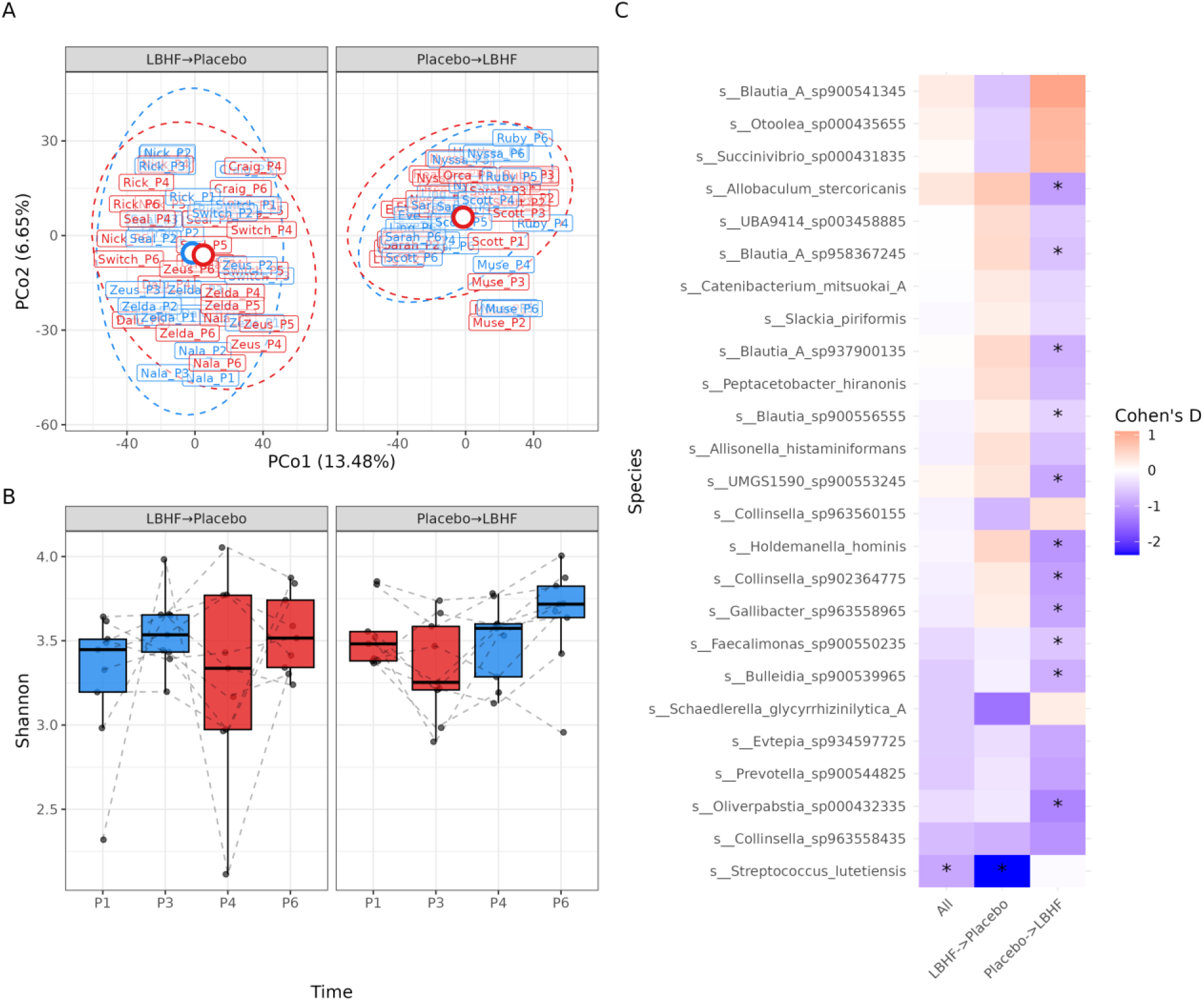
Gut microbiome composition and species-level responses across treatment sequences. **(A)** Principal coordinate analysis (PCoA) of Aitchison distances between species-level microbial profiles, stratified by treatment sequence (LBHF→placebo and placebo→LBHF). Samples are colored according to treatment. **(B)** Shannon diversity across sampling time points P1, P3, P4, and P6, stratified by treatment sequence. Boxplots show the distribution of Shannon diversity at each time point, and dotted lines connect repeated measurements from the same cat. Blue and red indicate LBHF and placebo treatment, respectively. **(C)** Species-level treatment-associated differences in within-cat changes in centered log-ratio (CLR)-transformed abundance, shown for the combined analysis and separately for the LBHF→placebo and placebo→LBHF sequences. Color indicates Cohen’s d, with negative values indicating a more negative change during LBHF treatment relative to placebo and positive values indicating a more positive change. Asterisks indicate FDR < 0.05.

Across both treatment sequences, the change in Shannon diversity was greater during LBHF treatment than during placebo, but the difference was not statistically significant (P = 0.08). In the placebo→LBHF sequence, the change in Shannon diversity differed between the LBHF and placebo periods (P = 1.25 × 10⁻³), whereas no corresponding difference was observed in the LBHF→placebo sequence (P = 0.70; **Figure 4B**).

We next evaluated within-cat changes in CLR-transformed species abundances between the beginning and end of each treatment period (**Figure 4C**; **Table S2**). In the combined analysis, *Streptococcus lutetiensis* was the only species with a significant treatment-associated difference after multiple-testing correction, showing a more negative change during LBHF treatment than during placebo (Cohen’s d = −0.90, FDR = 0.029). In the LBHF→placebo sequence, *S. lutetiensis* was also the only significant species (Cohen’s d = −2.37, FDR = 0.046). In the placebo→LBHF sequence, 11 species showed significantly more negative changes during the LBHF period than during the placebo period (FDR < 0.05; **Figure 4C**; **Table S2**).

As a sensitivity analysis, we restricted the comparison to Period A, thereby avoiding potential carryover effects. Changes in species abundance were compared between the cats receiving LBHF and those receiving placebo using Wilcoxon rank-sum tests (**Table S3**). Several taxa identified in the primary analysis showed nominal significance (P < 0.05) or showed trends towards significance (P < 0.10), including: *Streptococcus lutetiensis* (P = 0.05) and *Blautia* A sp937900135 (P = 0.04). However, none of these results met an FDR-adjusted significance threshold.

### LBHF-associated metabolomic changes differed between plasma and feces and by treatment sequence

Targeted metabolomic profiling detected 389 metabolites in fecal samples and 467 in plasma samples. Following prevalence filtering, 303 fecal and 431 plasma metabolites were retained for univariate analyses. At the global profile level, treatment accounted for little variation in either the fecal (PERMANOVA R² = 0.009, *P* = 0.56) or plasma (R² = 0.0086, *P* = 0.29) metabolome. Similarly, changes in metabolomic Shannon diversity did not differ significantly between LBHF and placebo in the overall analysis (fecal, *P* = 0.76; plasma, *P* = 0.15). In sequence-stratified analyses, the treatment-associated change in plasma Shannon diversity differed in the LBHF→placebo sequence (*P* = 0.01), with greater diversity during LBHF treatment, whereas no significant difference was detected in the placebo→LBHF sequence. No significant treatment-associated differences in fecal Shannon diversity were detected in either sequence (**Figure 5A,B**).

**Figure 5.**
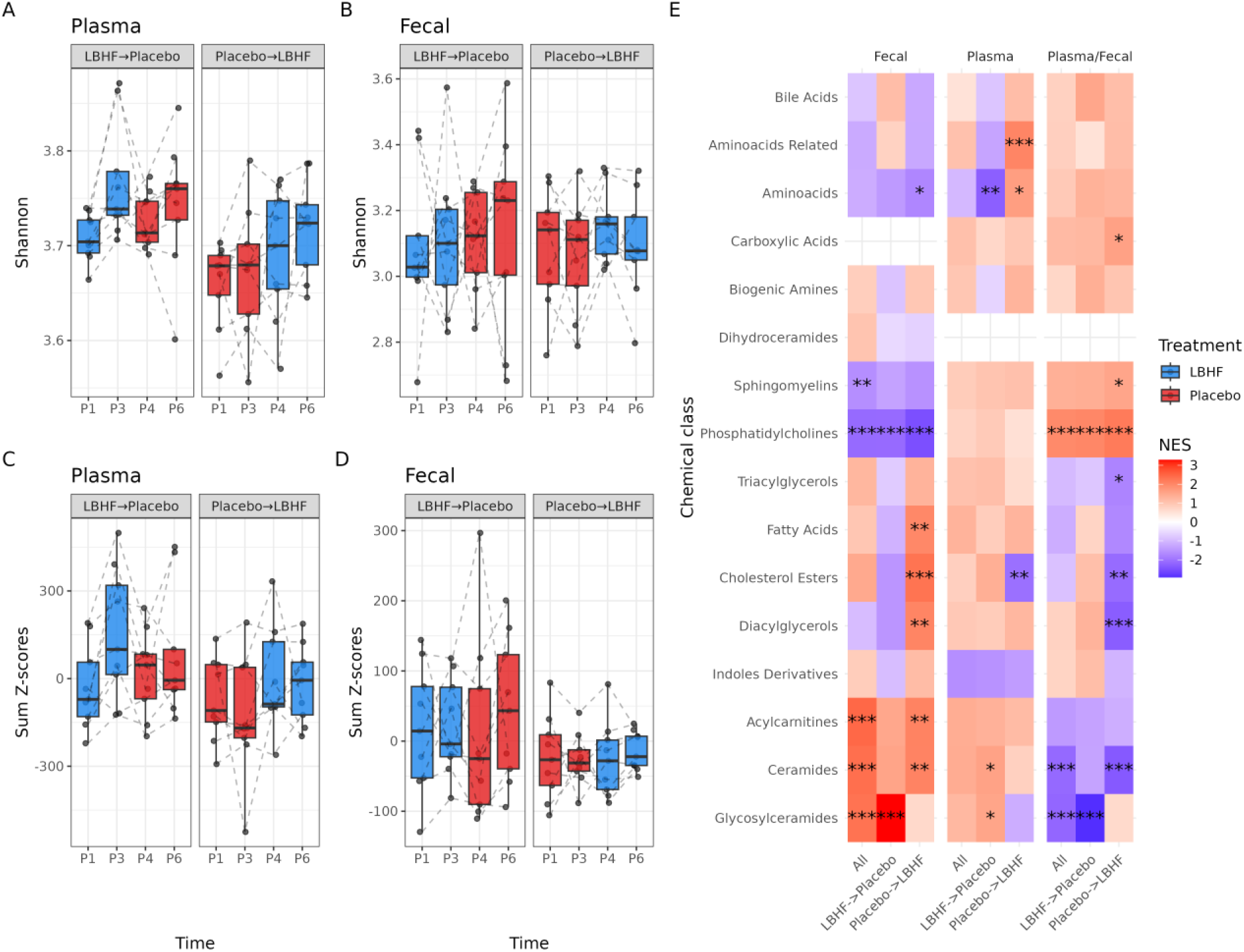
LBHF-associated changes in plasma and fecal metabolomic profiles. **(A, B)** Shannon diversity of plasma (A) and fecal (B) metabolomic profiles across study periods and treatment sequences. Each point represents an individual cat at a given sampling time (P1, P3, P4, or P6), with dotted lines connecting repeated measurements from the same cat. **(C, D)** Summed metabolite z-scores for plasma (C) and fecal (D) metabolomes across study periods and treatment sequences. Boxplots show the distribution of summed z-scores at each sampling point, and dotted lines connect repeated measurements from the same cat. Blue and red indicate LBHF and placebo treatment, respectively. **(E)** Metabolite-class enrichment associated with LBHF treatment across fecal, plasma, and plasma-to-fecal metabolite analyses. NES, normalized enrichment score. ***FDR < 0.001; **FDR < 0.01; *FDR < 0.05.

At the individual-metabolite level, 29 of 303 fecal metabolites and 33 of 431 plasma metabolites showed significantly different changes between LBHF and placebo after correction for multiple testing (FDR < 0.05; **Tables S4** and **S5**). All 33 plasma metabolites showed more positive changes during LBHF treatment than during placebo. Plasma metabolite concentrations generally decreased over each treatment period, but these decreases were smaller during LBHF treatment. In contrast, 28 of the 29 significant fecal metabolites showed more negative changes during LBHF treatment relative to placebo.

No metabolite classes were significantly enriched among the plasma metabolite associations in the combined analysis (FDR > 0.2; **Table S6**). In feces, phosphatidylcholines (NES = −2.12, FDR = 6.47 × 10⁻¹⁷) and sphingomyelins (NES = −0.74, FDR = 2.72 × 10⁻³) were negatively enriched, whereas glycosylceramides (NES = 2.30, FDR = 1.15 × 10⁻^5^), ceramides (NES = 2.45, FDR = 3.9 × 10⁻⁵), and acylcarnitines (NES = 2.35, FDR = 1.15 × 10⁻^5^) were positively enriched (**Table S6**).

Metabolite-level associations differed substantially between treatment sequences. In the LBHF→placebo sequence, 72 of 431 plasma metabolites differed significantly between treatments, 69 of which showed more positive changes during LBHF treatment (**Table S5**). In contrast, no individual plasma metabolites reached FDR < 0.05 in the placebo→LBHF sequence. For fecal metabolites, the opposite pattern was observed: eight of 303 metabolites differed significantly between treatments in the placebo→LBHF sequence, whereas none reached FDR < 0.05 in the LBHF→placebo sequence (**Table S4**). Sequence-specific metabolite-class enrichment results are provided in **Table S6**.

Consistent with the directional pattern observed for individual plasma metabolites, the summed plasma metabolite z-score showed a significant treatment-by-day interaction (*P* = 0.01; **Figure 5C**), whereas no corresponding interaction was detected for fecal metabolites (*P* = 0.40; **Figure 5D**). Finally, 37 of 245 plasma-to-fecal metabolite ratios differed significantly between LBHF and placebo after correction for multiple testing (**Table S7**). Thirty-six showed more positive changes during LBHF treatment, whereas indole-3-propionic acid showed a more negative change (Cohen’s *d* = −0.50, FDR = 0.04). Phosphatidylcholine plasma-to-fecal ratios were positively enriched during LBHF treatment (NES = 1.97, FDR = 1.21 × 10⁻¹¹; **Figure 5E**; **Table S6**).

### Digestibility was associated with distinct plasma and fecal metabolomic features

We next examined associations between digestibility endpoints and matched microbiome and metabolomic features. No species-level microbial features were associated with digestibility after correction for multiple testing, whereas several plasma and fecal metabolite associations were detected (**Table 3**). Plasma metabolites accounted for most of these associations, particularly for fat digestibility. Plasma CE(20:5) was positively associated with fat digestibility (FDR = 0.03), with additional positive associations with gross energy digestibility and metabolizable energy (FDR ≤ 0.10). Several additional plasma lipid metabolites were positively associated with fat digestibility, whereas plasma taurochenodeoxycholic acid (TCDCA) showed a negative association (**Table 3**). In feces, PC aa C32:3 was positively associated with gross energy digestibility and metabolizable energy (FDR ≤ 0.10). No plasma-to-fecal metabolite log-ratios were associated with digestibility after correction for multiple testing.

**Table 3.** Association between digestibility endpoint and metabolite level in plasma or fecal samples. Only associations with FDR ≤ 0.1 are shown.

| Source | Endpoint | Metabolite | Estimate | SE | P | FDR |
|---|---|---|---|---|---|---|
| Plasma | Fat | CE(20:5) | 19.69 | 4.05 | 5.54 × 10<sup>−5</sup> | 0.03 |
| Plasma | ME | CE(20:5) | 97.87 | 22.02 | 1.62 × 10<sup>−4</sup> | 0.08 |
| Plasma | Gross energy | CE(20:5) | 10.41 | 2.35 | 1.68 × 10<sup>−4</sup> | 0.08 |
| Fecal | ME | PC aa C32:3 | 32.81 | 6.82 | 2.06 × 10<sup>−4</sup> | 0.08 |
| Plasma | PC1 | CE(20:5) | 3.11 | 0.73 | 2.07 × 10<sup>−4</sup> | 0.10 |
| Fecal | Gross energy | PC aa C32:3 | 3.49 | 0.73 | 2.20 × 10<sup>−4</sup> | 0.08 |
| Plasma | Fat | TG(16:0_38:7) | 18.74 | 4.36 | 2.31 × 10<sup>−4</sup> | 0.05 |
| Plasma | Fat | lysoPC a C18:2 | 23.04 | 5.42 | 3.74 × 10<sup>−4</sup> | 0.06 |
| Plasma | Fat | Cer(d18:1/16:0) | 31.99 | 8.74 | 1.00 × 10<sup>−3</sup> | 0.10 |
| Plasma | Fat | CE(18:3) | 16.78 | 4.48 | 1.25 × 10<sup>−3</sup> | 0.10 |
| Plasma | Fat | TCDCA | −17.92 | 4.77 | 1.35 × 10<sup>−3</sup> | 0.10 |
| Plasma | Fat | HexCer(d18:1/23:0) | 20.23 | 5.71 | 1.44 × 10<sup>−3</sup> | 0.10 |
| Plasma | Fat | lysoPC a C20:4 | 19.55 | 5.40 | 1.66 × 10<sup>−3</sup> | 0.10 |

Class-level enrichment analysis further identified recurring lipid-related associations across digestibility endpoints (**Figure 6**). Across endpoints, enrichment profiles highlighted recurring lipid-related signatures, with phosphatidylcholines enriched among positively associated plasma metabolites and bile acid-related metabolites showing depletion signatures in both plasma and fecal analyses.

**Figure 6.**
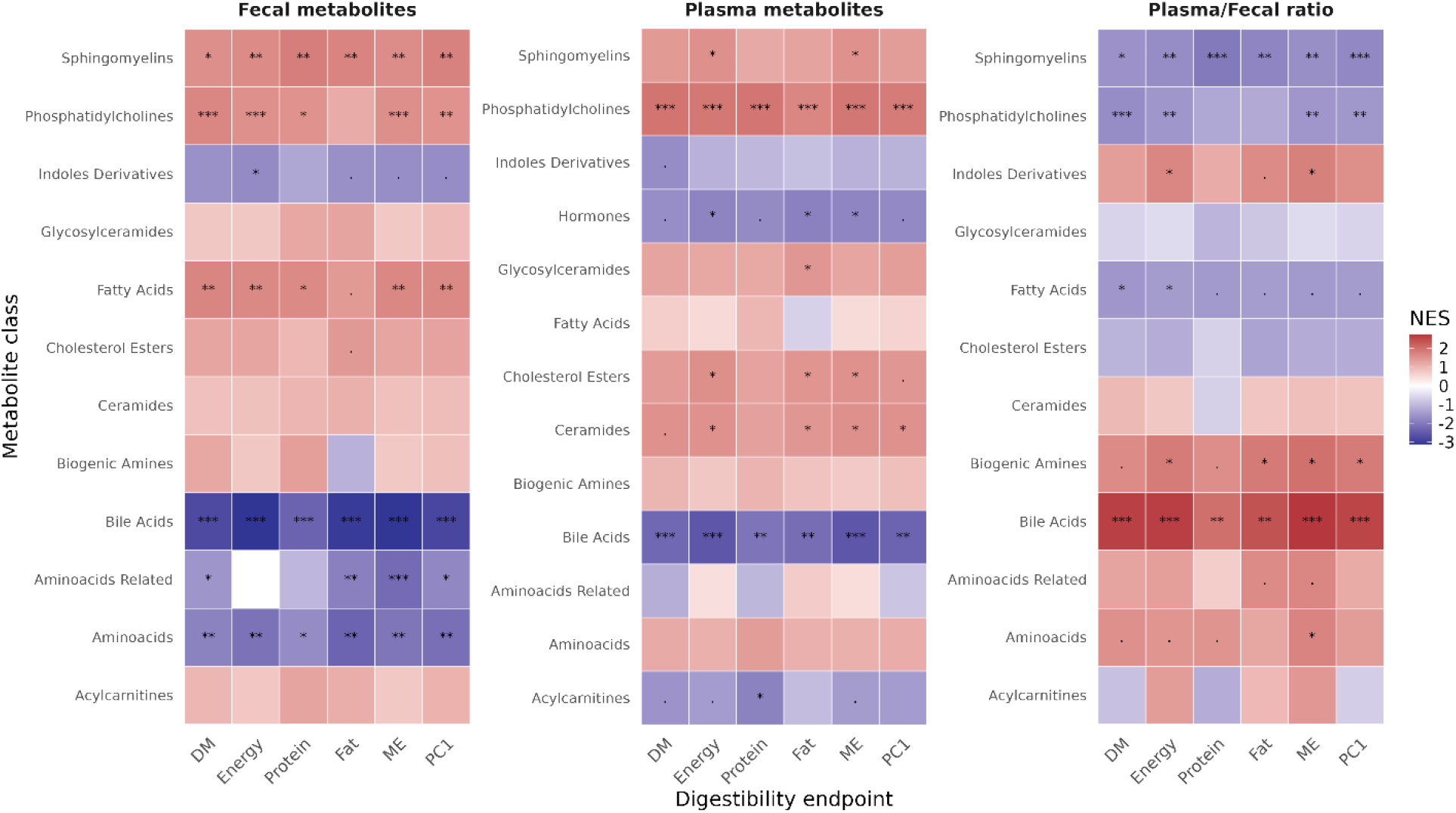
Metabolite-class associations with nutrient digestibility. Metabolite-class enrichment analyses were performed separately for plasma metabolites, fecal metabolites, and plasma-to-fecal metabolite log-ratios across apparent total tract digestibility endpoints. Metabolites were ranked according to the direction and statistical strength of their associations with each digestibility endpoint. Positive normalized enrichment scores (NES) indicate enrichment among metabolites positively associated with the corresponding digestibility endpoint, whereas negative NES values indicate enrichment among negatively associated metabolites. Only associations meeting the displayed FDR thresholds are indicated. ***FDR < 0.001; **FDR < 0.01; *FDR < 0.05.

## Discussion

In this placebo-controlled crossover study of older cats, LBHF supplementation did not affect intestinal permeability, the primary endpoint, but was associated with improved nutrient digestibility, with the clearest endpoint-specific effect observed for protein digestibility. LBHF was also associated with changes in the plasma and fecal metabolomes and more modest changes in the gut microbiome, although several of these responses differed markedly by treatment sequence. Together, these findings suggest that the effects of LBHF in older cats may relate more strongly to digestive efficiency and metabolic responses than to modulation of intestinal permeability.

The primary objective of this study was to test if LBHF could mitigate age-related increases in gut barrier permeability in a colony of aged cats. Previous studies in *C. elegans* (Eckhardt et al., manuscript in preparation) and human colonic 3D tubuloid explants had shown that LBHF was able to reduce gut barrier leakiness (Cercamondi et al., 2025). Although LBHF did not reduce intestinal permeability compared with placebo, we also observed the previously reported positive association between age and intestinal permeability in this colony of cats (Patterson et al., 2026). The lack of treatment effect may reflect several factors, including limited statistical power, variability in permeability measurements, treatment duration, or the relatively low number of very old animals exhibiting the highest permeability values.

Alternatively, gut leakiness across an otherwise intact gut barrier may not be the right marker to assess gut barrier function. Epithelial damage beyond tight junction leakiness, such as enterocyte necrosis or sloughing of sections of epithelia due to microbial or toxicological challenges, might be more suitable for detecting possible potential postbiotic effects. The present study did not measure any marker indicative of these phenomena. However, studies on weaned pigs showed that a challenge of the upper gastrointestinal tract with enterotoxigenic *E*. *coli* led to less jejunal mucosal stress when pigs were fed with LBHF compared to controls (Xu et al., 2022; Duarte et al., 2024). LBHF previously also appeared to reduce intestinal epithelial burden in mice caused by infections with *Citrobacter rodentium*, *Campylobacter jejuni*, or *Salmonella enterica* serovar Typhimurium (Moyen et al., 1986; Coconnier et al., 1997; Warda et al., 2020). These observations suggest that the gut barrier protection mechanisms of LBHF may be more related to fending off challenges than tightening the junctions between epithelial cells.

The most consistent physiological response to LBHF was improved nutrient digestibility, with the strongest endpoint-specific evidence observed for protein digestibility. Because protein digestion and absorption occur predominantly before the large intestine, this finding suggests that LBHF may influence digestive processes in the upper gastrointestinal tract. Previous *in vitro* studies have reported effects of Lactobacillus-derived preparations on brush-border enzymes involved in nutrient digestion and absorption, including alkaline phosphatase, dipeptidyl peptidase IV, and sucrase-isomaltase (Liévin-Le Moal et al., 2002). Whether similar mechanisms contributed to the digestibility effects observed here remains to be established.

Improved protein digestibility may be especially relevant in older cats, for whom maintaining adequate protein uptake is important for preservation of lean body mass and healthy aging. Furthermore, greater digestion of dietary protein in the small intestine may reduce protein spillover to the colon, limiting substrate availability for proteolytic fermentation and the production of potentially detrimental microbial metabolites (Badri et al., 2021; Summers et al., 2023). Digestibility studies comparing diets with strong differences in protein levels facilitated detection of significant changes in microbial composition and function in the hindgut (Badri et al., 2021; Summers et al., 2023). However, the present study did not directly quantify protein delivery to the colon, and the observed microbiome changes were modest. Consequently, whether altered substrate availability contributed to the microbial or metabolomic responses remains unknown.

We observed several associations between ATTD endpoints and plasma and fecal metabolite changes. Notably, cholesteryl eicosapentaenoate 20:5 (CE(20:5)) was associated with increased fat and energy digestibility and metabolizable energy. Eicosapentaenoic acid (EPA) plasma levels have previously been linked to anti-inflammatory effects and are associated with a decrease in cardiovascular mortality and all-cause mortality in humans (Miura et al., 2016; Leleu et al., 2025). In companion animals, EPA supplementation has demonstrated benefits for many indications (Magalhães et al., 2021). Associations of EPA with increased fat and energy digestibility and metabolizable energy in our study could mean that increased digestibility upon LBHF treatment could improve overall lipid digestibility and EPA bioavailability, although it does not establish a causal role of LBHF on lipid absorption or transport.

Significant microbiome and fecal metabolome responses were observed primarily in the placebo→LBHF sequence, suggesting a potential treatment-order effect. One plausible explanation is an incomplete washout between intervention periods. Previous studies have shown that microbiome responses to microbial interventions can persist for weeks after treatment cessation, potentially obscuring subsequent treatment effects (Whittemore et al., 2019). Consistent with this interpretation, Shannon diversity increased significantly during LBHF administration in the placebo→LBHF sequence but not in the LBHF→placebo sequence, suggesting that the microbiome may not have fully returned to baseline before the crossover. Future studies incorporating longer washout periods may be better positioned to distinguish transient from persistent LBHF-associated effects. Additionally, long-lived LBHF effects might also be studied with longer follow-ups after intervention to assess long-term benefits of LBHF in both microbiome and host physiology.

Interestingly, the reduction in *S. lutetiensis* abundance may also be viewed from a One Health perspective. Although the significance of this organism in feline health remains poorly understood, in humans the bacterium is considered an emerging pathogen and has been isolated from blood culture during bacteremia, from dairy cows with mastitis, and from tumor tissues of gastric cancer patients (Almuzara et al., 2013; Chen et al., 2021; Kodilinye et al., 2023). Mouse models confirm that the bacterium promotes cancer progression, but its role in cats is largely unknown and limited to its reported isolation from a diseased cat (Piva et al., 2019; Wang et al., 2025). Members of the gut microbiota can be transmitted between companion animals and humans through close household contact (Heidrich et al., 2025). Consequently, interventions that reduce the abundance of potential pathobionts in pets may have implications beyond the individual animal, potentially influencing microbial exchange within shared human-animal environments.

The predominantly more positive plasma metabolite changes observed during LBHF treatment could reflect differences in intestinal absorption, microbial metabolism, host metabolism, tissue uptake, or clearance. The present study cannot distinguish among these mechanisms. Similarly, changes in plasma-to-fecal metabolite ratios should not be interpreted as direct measures of intestinal absorption because both compartments are influenced by multiple metabolic processes. Stable-isotope tracing and targeted mechanistic experiments could help determine whether the observed systemic metabolomic differences reflect altered nutrient absorption or downstream metabolic responses.

In conclusion, LBHF was well tolerated in older cats and produced measurable physiological effects despite failing to improve intestinal permeability, the primary endpoint of the study. We observed an improvement in nutrient digestibility, particularly protein digestibility, accompanied by broad changes in the plasma metabolome and more modest alterations in the gut microbiome and fecal metabolome. These findings suggest that the principal benefits of LBHF in healthy older cats may relate more to digestive efficiency and host metabolic responses than to direct modulation of gut permeability. The metabolomic signatures identified here provide several hypotheses for future mechanistic studies and support further evaluation of postbiotic interventions in aging companion animals. Our study provides a proof-of-principle and framework for future studies in other cat populations to evaluate the effects of postbiotic interventions on intestinal permeability, the microbiome, and the metabolome.

## Acknowledgements

The authors thank Jennifer Tsang, LLC, for providing writing and editorial support and Rosa Argamasilla for support with the metabolomic measurements.

## Author Contributions

E.E. – Review and editing, conceptualization, investigation, project administration, supervision

S. A.-S. – Methodology, formal analysis, writing original draft, review and editing

B.T. – Methodology, formal analysis, review and editing

A.-S.B. – Methodology, investigation

B.K. – Methodology, investigation, project administration, review and editing

E.P. – Conceptualization, project administration

K.F. – Conceptualization, project administration, formal analysis, review and editing

M.A.v.d.B – Review and editing, supervision

D.T. – Conceptualization, project administration, review and editing

M.H. – Methodology, study execution and supervision, review and editing

D.R.B. – Formal analysis, review and editing

## Ethics approval declaration

The protocol for this study was approved by Massey University Animal Ethics Committee: AEC 24/36, granted 13 September 2025. All cats used for this study were housed at the Centre for Feline Nutrition, Massey University, Palmerston North, New Zealand according to the Animal Welfare (Companion Cats) Code of Welfare (2007).

## Abbreviations

CLR: centered log-ratio
ESI: electrospray ionization
FDR: false discovery rate
GSEA: gene set enrichment analysis
LC-MS: liquid chromatography-mass spectrometry
L:R: lactulose:rhamnose ratio
PC1: first principal component
PERMANOVA: permutational multivariate analysis of variance
QC: quality control
UHPLC: ultra-high-performance liquid chromatography
X:G: xylose:3-O-methylglucose ratio

## Data availability

Raw metagenomic sequencing reads generated in this study have been deposited in the NCBI Sequence Read Archive under BioProject accession PRJNA1489369.

## Supplemental Material

### Supplemental Figures

**Supplementary Figure 1.**
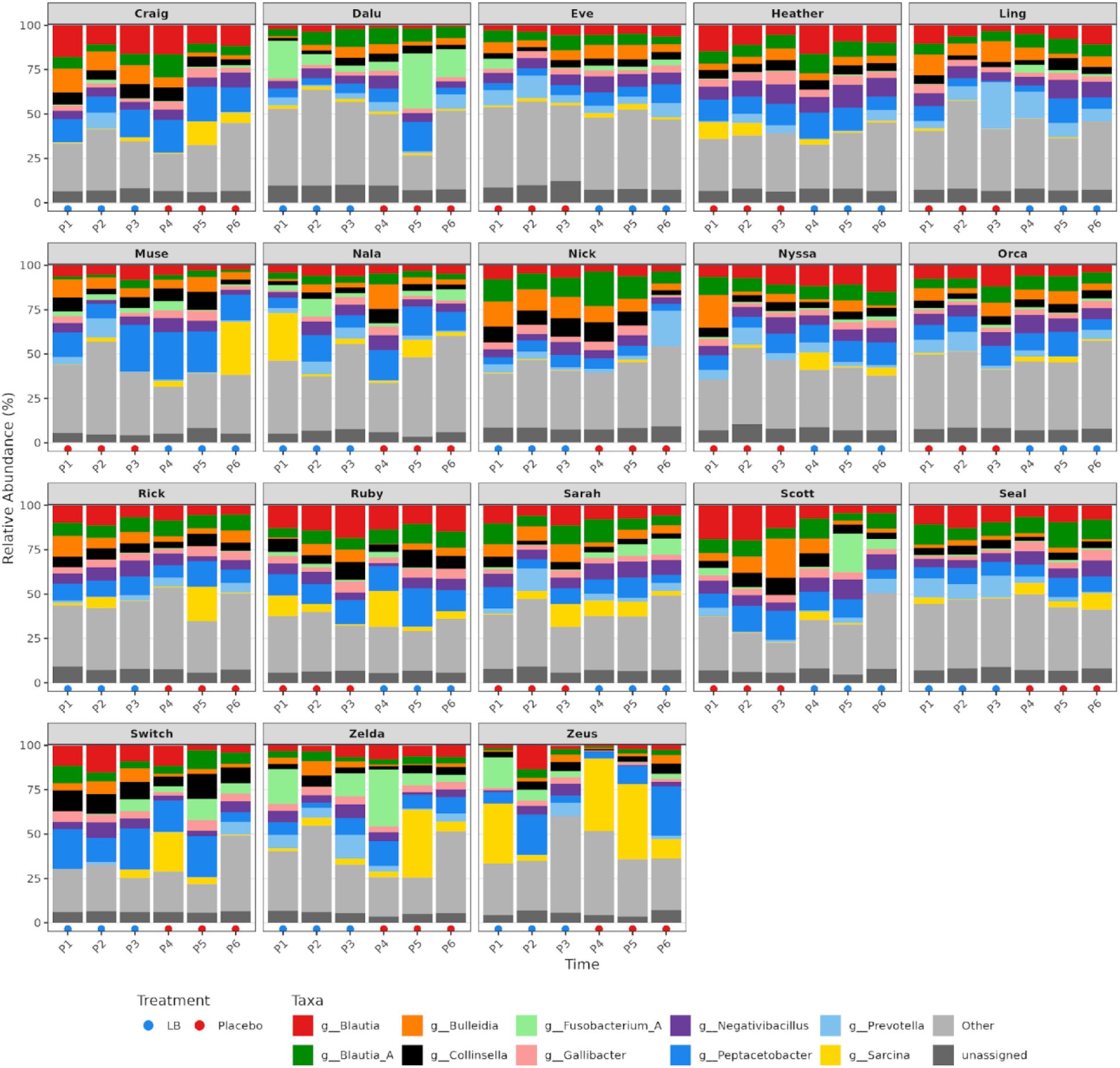
Genus-level composition of feline gut microbiomes across the study. Stacked bar plots show the relative abundance of microbial genera for fecal samples from each individual cat across six experimental time points (P1–P6). Each panel represents one animal. Colors indicate the relative contribution of each genus to the total community composition, while “Other” captures low-abundance genera not individually displayed, and “Unassigned” denotes fractions without a genus-level taxonomic assignment. Treatment at each time point is indicated by point color along the x-axis (blue = LBHF; red = placebo). Across all time points from both treatment sequences, the feline gut microbiome was dominated by Bacillota (median relative abundance: 80.5%), followed by Bacteroidota (6%) and Actinomycetota (5.5%). At the genus level, Peptacetobacter was most abundant (median 9.5%), with Blautia A (6.9%), Blautia (6.7%), Negativibacillus (5.9%), and Collinsella (4.7%) also contributing substantially to community composition.

