## Supplementary Figure 1 for "Effect of *Lactobacillus* postbiotic supplementation on intestinal permeability, nutrient digestibility, and gut microbial and metabolic profiles in aging cats"

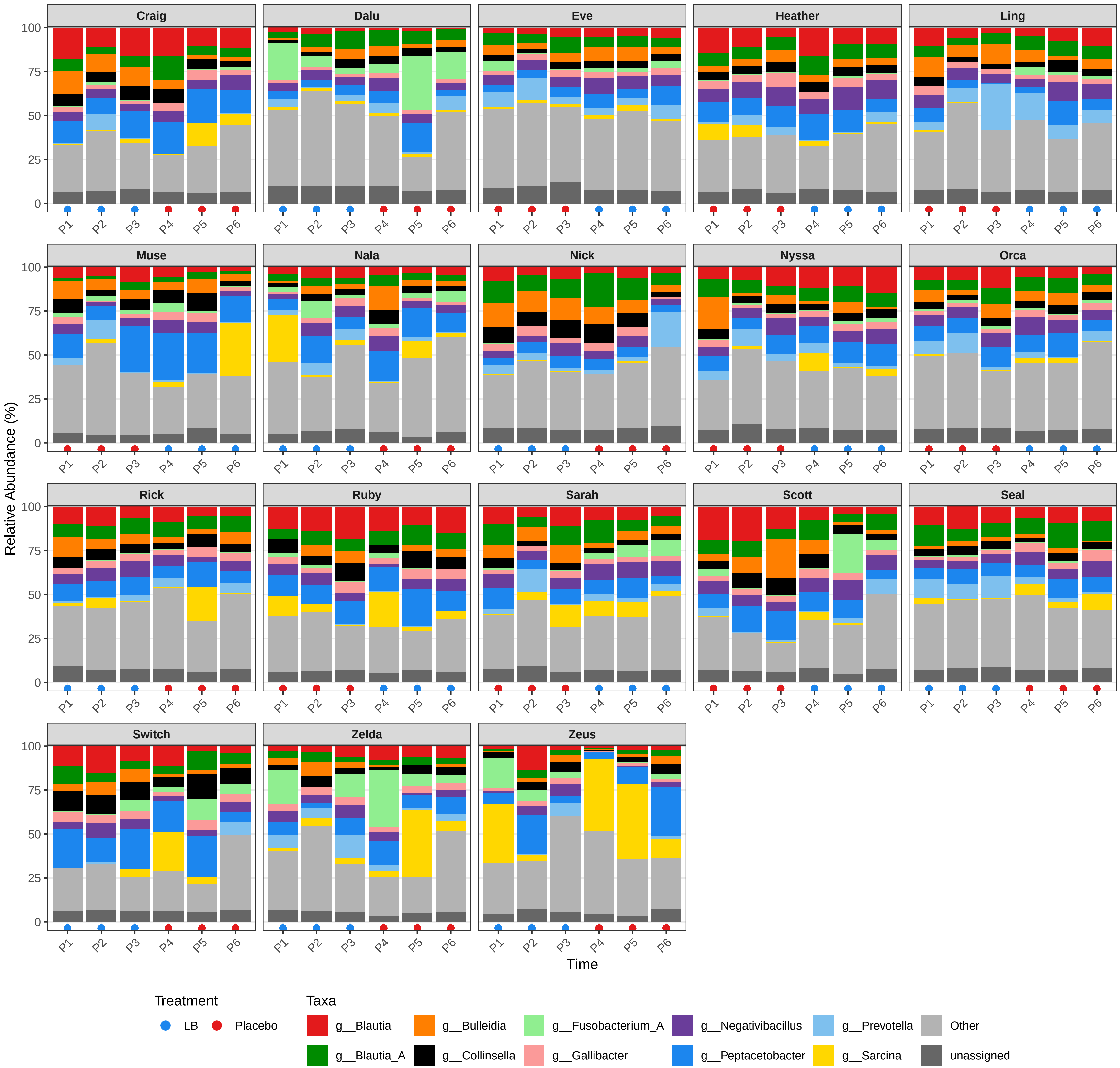

Craig
Dalu
Eve
Heather
Ling
Muse
Nala
Nick
Nyssa
Orca
Rick
Ruby
Sarah
Scott
Seal
Switch
Zelda
Zeus
Relative Abundance (%)
0
25
50
75
100
P1
P2
P3
P4
P5
P6
Time
Treatment
LB
Placebo
Taxa
g__Blautia
g__Blautia_A
g__Bulleidia
g__Collinsella
g__Fusobacterium_A
g__Gallibacter
g__Negativibacillus
g__Peptacetobacter
g__Prevotella
g__Sarcina
Other
unassigned
